# Joint seasonality in geographic and ecological spaces, illustrated with a partially migratory bird

**DOI:** 10.1101/514349

**Authors:** Mathieu Basille, James Watling, Stephanie Romañach, Rena Borkhataria

## Abstract

As most species live in seasonal environments, considering varying conditions is essential to understand species dynamics in both geographic and ecological spaces. Both resident and migratory species need to contend with seasonality, and balance settling in favorable areas with tracking favorable environmental conditions during the year. We present an exploratory framework to jointly investigate a species’ niche in geographic and ecological spaces, applied to wood storks (*Mycteria americana*), which are partially migratory wading birds, in the southeastern U.S. We concurrently described monthly geographic distributions and climatic niches based on temperature and precipitation. Geographic distributions of wood storks were more similar throughout the year than were climatic niches, suggesting that birds stay within specific areas seasonally, rather than tracking areas of similar climate. However, wood storks expressed consistent selection of warm areas during the winter, and wet areas during the summer, indicating that the selection of seasonal ranges may be directly related to environmental conditions across the entire range. Our flexible framework, which simultaneously considered geographic and ecological spaces, suggested that tracking climate alone did not explain seasonal distributions of wood storks in breeding and non-breeding areas.

## Introduction

At the core of ecology is the question of where do animals live? Early on, ecologists acknowledged the dual nature of this issue, by investigating species ranges (i.e., their location in geographic space, Candolle 1855, Darwin 1859, Wallace 1876), and species ecological niches (i.e., their location in ecological space, Grinnell 1917, Hutchinson 1957), actually reflecting what has been termed “Hutchinson’s duality” (Colwell and Rangel 2009). More recent is the joint investigation of the effect of a change in one space on the other—e.g., how range constraints caused by migration or breeding affect a species’ niche (Kearney and Porter 2009), or how climate change may affect species’ range limits, (Chen et al. 2011). However, the dynamic nature of both spaces in time—i.e., their *seasonality*—often goes unacknowledged, despite the fact that most species live in seasonal environments (Fretwell 1972), with strongly cyclic variations in resource availability and conditions throughout the year. Here we illustrate the use of a generalized framework for joint seasonality in geographic and ecological (specifically climatic) spaces, with the case of wood storks (*Mycteria americana*) in the southeastern United States.

The direct correspondence between environmental conditions (i.e., ecological niche) and a species range at equilibrium is the fundamental assumption of species distribution modeling (Araújo and Pearson 2005, Pearman et al. 2008, Elith and Leathwick 2009). However, variations of environmental conditions during the year has led to the recognition of seasonal variation in ecological niches, with species responses varying on a continuum from “niche trackers” to “niche switchers” (Martínez–Meyer et al. 2004, Nakazawa et al. 2004). Broennimann et al. (2011) proposed a framework to quantify niche overlap using essentially kernel overlap metrics in a simplified 2-dimensional niche. Relying on this framework, several studies investigated seasonality in ecological niches: for instance, Laube et al. (2015) estimated overlap between breeding and non-breeding niches of warblers, while Gómez et al. (2016) investigated seasonal niche overlap of passerine birds.

Movement has evolved as the primary means to manage heterogeneous environments in space or in time (Nathan et al. 2008). In seasonal environments, motile animal species can be placed on a gradient from sedentary to migratory (Chapman et al. 2011, Cagnacci et al. 2011, Jonzén et al. 2011), with direct implications for seasonal overlap, both in geographic and ecological spaces. Alternatively, nomadism can be thought of as a response to highly variable and unpredictable environments in both time and space (Teitelbaum and Mueller 2019), which should lead to variable ranges through time, as species track their ecological niche closely (Jonzén et al. 2011). On the one hand, sedentary species should exhibit a high similarity in their range, but will experience environmental seasonality, resulting in low ecological similarity throughout the year. On the other hand, migration is a dramatic response to seasonal environmental change (Dingle and Drake 2007), which leads to two distinct seasonal ranges. As a consequence, occupied space should show high similarity within seasons in both geographic and ecological spaces, but not necessarily between seasons, with migrations potentially resulting in shifts in both spaces. Some species also migrate within their range, e.g., through altitudinal migration (i.e., migratory range shifts along an elevation gradient, Zweifel-Schielly et al. 2009). These species mostly stay within their global range year-round, while locally buffering for seasonally varying conditions to some extent. As a result, it is expected that their occupied space should vary throughout the year, but with intermediate patterns compared to the high expected similarity of residents in geographic space, and of migrants in ecological space.

Finally, it has been recognized that many species express a form of partial migration (a fraction of the population is migratory, while the other part remains resident, Chapman et al. 2011), or facultative migration (individuals that may or may not migrate in a given year, Newton 2012). Partial or facultative migrants pose a conundrum, as the global geographic range and environmental conditions experienced may encompass distinct and contrasted subsets (for residents and migrants) during part of the year or in different years. This plasticity allows facultative migrants to pick and choose when to migrate in response to environmental conditions or internal state, as opposed to presumably more hard-wired obligate migrants. In the case of partial migration, we thus expect the mix between the two strategies (resident or migrant) to be reflected in both geographic and ecological spaces, resulting, at the population level, in intermediate similarity between that of resident and migrant species during the migratory season.

Here we describe the seasonality in the distribution of wood storks, which are partially migratory birds (Coulter et al. 1999, Picardi et al. 2020),, in both geographic and ecological (specifically climatic) spaces. We built on the approaches advocated by Fieberg and Kochanny (2005) in the geographic space only, and by Broennimann et al. (2011) in the ecological space only, to propose a general framework that allows joint investigation in geographic and ecological spaces. To facilitate combined interpretation in both spaces, we directly used a simplified 2-D niche space with two critical climatic components varying throughout the year: temperature and precipitation. We use home range overlap metrics (Fieberg and Kochanny 2005) to simultaneously investigate similarity of monthly ranges in geographic space and monthly niches in climatic space. Because wood storks are partial migrants breeding in South Florida during winter (dry season, Coulter et al. 1999), we expect both monthly ranges and niches to be more restricted and highly similar during this season. On the other hand, as a fraction of the population migrates to northern foraging grounds after breeding, we then expect both monthly ranges and niches to show low similarity during the rest of the year.

## Methods

### Wood Storks in southeastern U.S

Sixty-one adult wood storks were captured between 2005 and 2011 from 11 sites spread throughout their southeastern U.S. range in Florida, Georgia, and South Carolina. Wood storks were equipped with solar-powered Argos/GPS tags set up to take a fix every 1h or 2h. Tagged birds were monitored for an average of 649 ± 563 d from July 2005 to December 2011, and the tags collected a total of 445,638 GPS locations, which is more than 7,000 locations per individual on average (7, 306 ± 6, 036). We defined the study area by the coastline to the South (i.e., all land masses), and by the limit of a convex hull around all GPS locations to the North (see the study area limits in Fig. 1).

**Figure 1:**
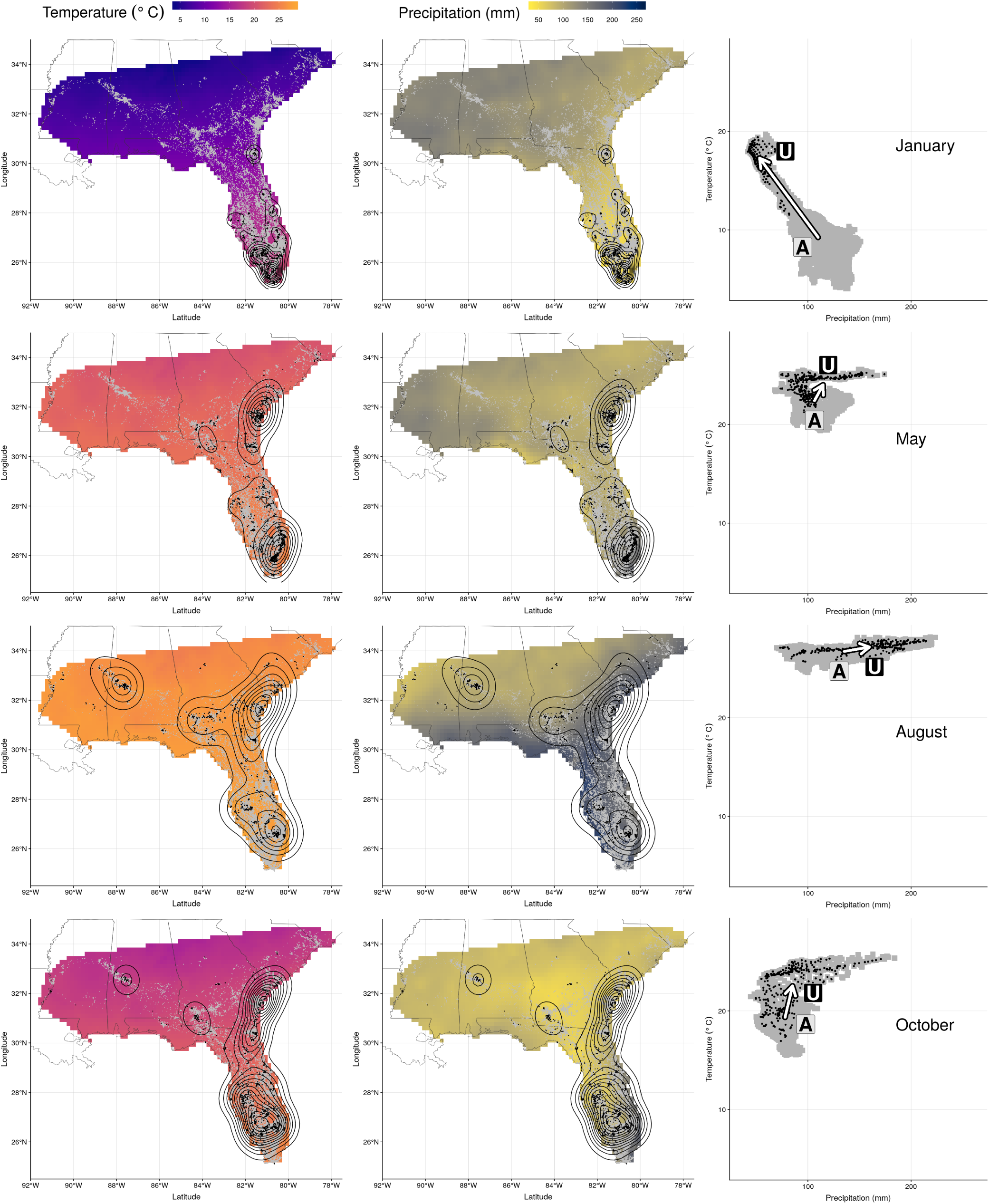
Wood stork monthly range (left and center) and climatic niche (right) during January, May, August and October. Range maps show current GPS locations in a given month in black with kernel contour lines, and all locations from the entire dataset shown as a reference in gray, on a background of temperature (left, from blue for colder temperatures to red for warmer temperatures) or precipitations (center, from yellow for dry conditions to blue for wet conditions). Climatic niches show current conditions used in a given month (precipitation on the X-axis, and temperature on the Y-axis) in black, in comparison to conditions available in the study area in a given month in gray. The marginality arrow defines the difference between average available conditions (“A”) and average used conditions (“U”).

We first grouped all GPS data by month across all years to define monthly ranges. To account for variability in number of locations among individuals, and the fact that individual wood storks are a random sample of the population, we sampled 5,000 GPS locations in each month by 1) removing individuals with fewer than 100 locations in a given month, and 2) sampling 5,000 random individual wood storks (with replacement) and 1 random location for each of the 5,000 sampled individuals. We thus compiled 60,000 random locations for the population, evenly distributed throughout the year.

In climatic space, the wood stork niche was described by average monthly temperature and precipitation using climate rasters from the Climate Research Unit (CRU) data set, available at a resolution of approximately 10 arc-min (New et al. 2002). A previous analysis (Watling et al. 2012) showed that variables describing monthly climate (e.g., mean monthly temperature and precipitation) performed equally well as the more ecologically sensible “bioclimate” variables derived from seasonal relationships between precipitation and temperature (e.g., maximum temperature of the warmest month, or mean precipitation of the driest month) to define climatic niches of twelve terrestrial vertebrate species in the southeastern U.S. Remarkably, the only exception was wood storks, which can be explained by the global range of the species spanning both hemispheres, hence effectively preventing monthly variables from be meaningful (Watling et al. 2012). Focusing on the southeastern U.S. range only circumvents this issue, and actually supports the use of monthly variables as the simplest approach.

Global layers were clipped to the study area, and temperature and precipitation at sampled locations defined a 2D-climatic niche, similar to the range defined in the 2D-geographic space (longitude and latitude) by GPS locations. To allow meaningful comparison between temperature/precipitation and longitude/latitude, the former were standardized as to be on the same scale (subtraction of the mean followed by division by the standard deviation for all locations).

### A note on ecological niches

At this stage, it is important to note two things: First of all, as in every species distribution model, what we observe by means of occurrence data does only reflect realized niches (Colwell and Rangel 2009, Soberón and Nakamura 2009), i.e., realizations of the fundamental niche of a species in a given context, restricted by biotic interactions and movement or dispersal constraints (Soberón 2007). Several studies have tried to circumvent this limitation (Broennimann and Guisan 2008, Panzacchi et al. 2015), but they require several populations in different ranges to approach the fundamental niche. In this manuscript, we thus only consider the realized niche through the year. Niches can change through time, and such changes observed in the realized niche also include any potential change of the fundamental niche, although it is impossible to disentangle both (Guisan et al. 2014).

Second, the seminal work of Hutchinson (1957) provided at the same time a conceptual framework to study both fundamental and realized niches, and a clear geometrical model of the niche, defined as a “*n*-dimensional hypervolume”, i.e., a multivariate cloud of points in the ecological space. This model can be useful to investigate niches, whether we consider fundamental, potential, or realized niches (Soberón and Nakamura 2009). As a matter of fact, we rely on this model to evaluate how the niche changes through time, and we extend it to define the concept of monthly niche: just as we can measure “monthly ranges”, defined from all geographical locations in a given month, we define “monthly niches”, as the realization of the fundamental niche in a given month—i.e., a realized niche at a monthly scale.

### Geographic and climatic similarity

Our framework relies on the observation that the problems of measuring home range overlap in the geographic space (Fieberg and Kochanny 2005), and measuring niche overlap in the ecological space reduced to a 2D Cartesian plane (Broennimann et al. 2011) are actually mathematically identical problems, which reduce to defining what metric of overlap to use. In the context of home ranges, Fieberg and Kochanny (2005) recommend Bhattacharyya’s affinity (BA), a synthetic measure (i.e., symmetric) appropriate to quantify the overall similarity of two bivariate probability densities, and ranges from zero (no overlap) to 100% (complete overlap). In the context of niche overlap, Broennimann et al. (2011) presented a method to compute overlap between two ecological niches (i.e., in the ecological space) in three steps: 1) reducing the multidimensional space of the niche to two dimensions; 2) compute kernel densities to circumvent potential sampling biases; 3) estimate overlap using Schoener’s *D*. Warren et al. (2008) however recommend the use of a derivative of Hellinger’s distance over Schoener’s *D*. It can actually be shown that their newly presented metric 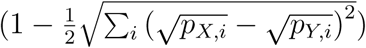 corresponds mathematically to BA, which provides fundamental justification for the use of BA as an appropriate metric of overlap.

We thus computed utilization distributions (UDs), i.e., the bivariate probability density associated with wood storks in both geographic (defined by longitude and latitude) and climatic (defined by standardized temperature and precipitation) spaces using the kernel method (Worton 1989) with standard parameterization (*ad-hoc* method for the smoothing parameter, which supposes a bivariate normal UD). We then measured overlap in both geographic and climatic spaces throughout the year with Bhattacharyya’s affinity, which defines seasonal similarity (Cagnacci et al. 2016).

### Marginality

Different constraints acting in the climatic and geographic spaces render quantitative comparison of overlap in both spaces difficult. Instead, as the geographic range bounds what is available to a species, it is possible to better understand the dynamics at play in climatic space by further investigating monthly selection of climatic conditions by means of marginality. Marginality, the vector connecting the centroid of environmental conditions in the geographic area and the centroid of conditions in the realized niche, measures the difference between available and used conditions (Hirzel et al. 2002, Basille et al. 2008). In our case, we used marginality to measure the difference between climatic conditions in the study area and climatic conditions at wood stork GPS locations as temperature and precipitation change month by month throughout a year. Note here that the study area, in climatic space, is completely defined throughout the year as the union of the 12 monthly climate layers.

All analyses were performed in R (R Core Team 2017), with the help of the “adehabitatLT” package (Calenge 2006) to process GPS locations. All scripts are available in a GitHub repository (https://basille.github.io/wost-seasonality/).

## Results

There was greater similarity (as measured by overlap) in geographical space than in climatic space (Fig. 2), despite large geographic variation in wood stork distribution during the year (see the full video in Metadata S1). Overall, similarity in the geographic range between any two months was always greater than 50 %, with minima reached between the months of January–February and July–September (consistently between 51–55 %). On the other hand, similarity in the climatic space was much lower, ranging from 0 % between many months throughout the year, to a maximum of 83 % between July and August. In particular, similarity from successive months ranges from 6 % to 83 % (mean±SD: 42.0±23.5) in the climatic space vs. 89 % to 99 % (mean±SD: 94.8±3.4) in the geographic space.

**Figure 2:**
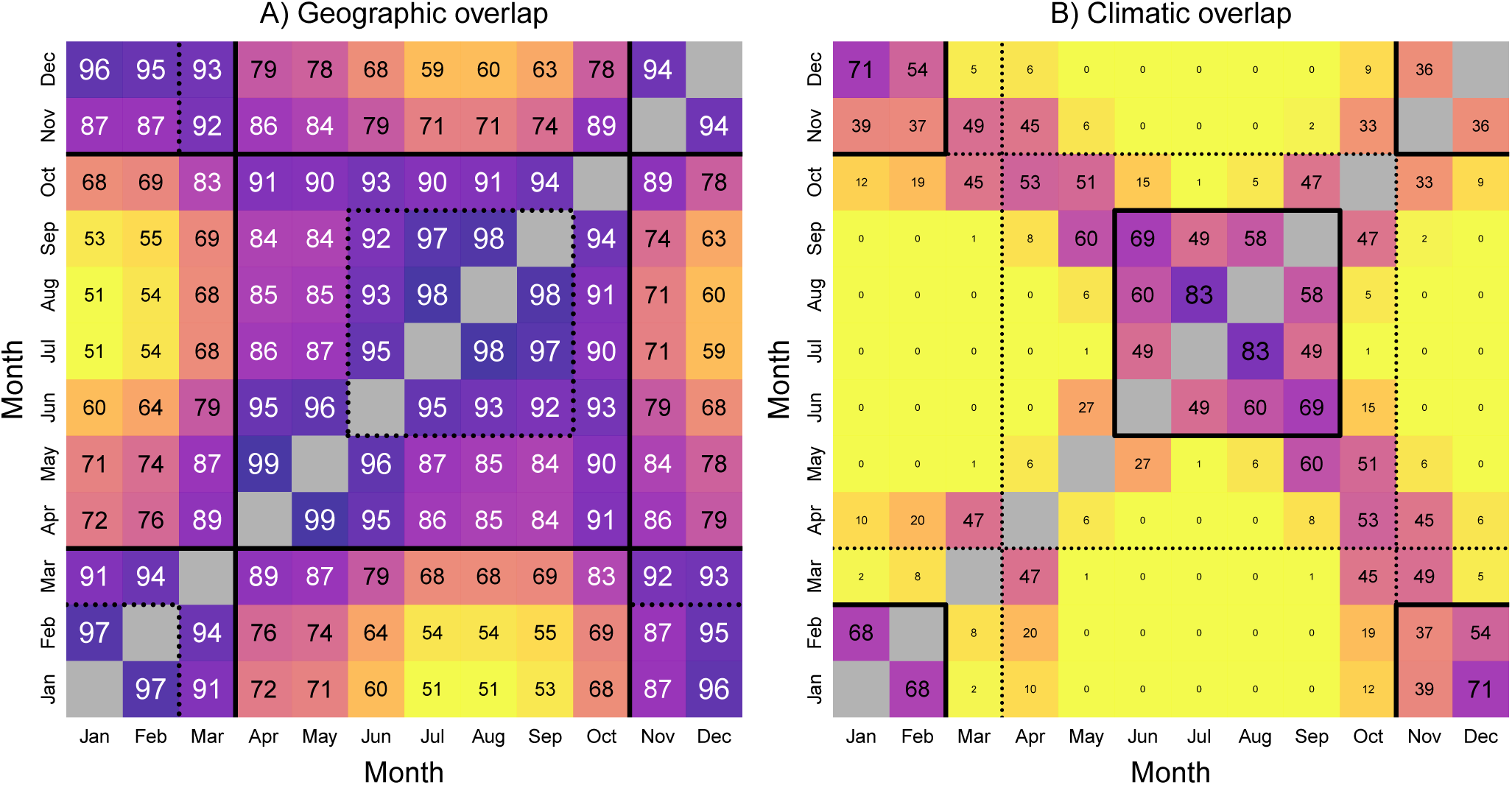
Matrices of A) geographic and B) climatic overlap during the year. Each matrix provides Bhattacharyya’s affinity as a percentage (between 0 and 100, higher values corresponding to darker color), as a measure of overlap between two months *i* (rows) and *j* (columns). The main diagonal, where *i* = *j* is grayed out as all values equal 1. Consistent seasons in a given space are highlighted with a bold line, while seasons identified in the other space are highlighted with a dotted line for reference.

The different patterns of seasonal change in geographic and climatic space confound interpretation of the overlap in both spaces. For instance, the use of a high threshold in the geographic space (e.g. 80 %) delineates two very consistent seasons, Summer from April to October included, and Winter from November to March included (Fig. 2). However, transitions from each season are very smooth, with high values of overlap (89 %) in both cases. In other words, the change of the geographic distribution is very progressive from month to month. Conversely, using thresholds as low as 35 % in the climatic space still delineates two seasons, Summer from June to September included, and Winter from November to February included, which are actually subset of the seasons previously defined in the geographic space (Fig. 2).

Within-season similarity is greater in geographic space than in climatic space: the lowest similarity between any two months for a given season is 84 % (between April and September, and between May and September), while similarity drops to 1 % in the climatic space, even in the subset seasons (between July and October) (Fig. 2). This highlights both the high similarity of monthly ranges as compared to monthly climatic niches, but also the higher discriminatory power of the latter, due to larger contrasts.

Finally, monthly marginality in climatic space highlighted a very clear picture of an annual cycle (Fig. 3): wood storks consistently used areas wetter and slightly warmer than generally available in their range during the summer (July–August), and consistently used warmer and drier areas during the winter (November–April). Transitional months between these two seasons appeared clearly to be May and October, with areas used by wood storks in these two months warmer and slightly wetter than generally available in their entire range. In fact, marginality in May and October were very similar, and much more similar to each other than to any other month of the year (Fig. 3).

**Figure 3:**
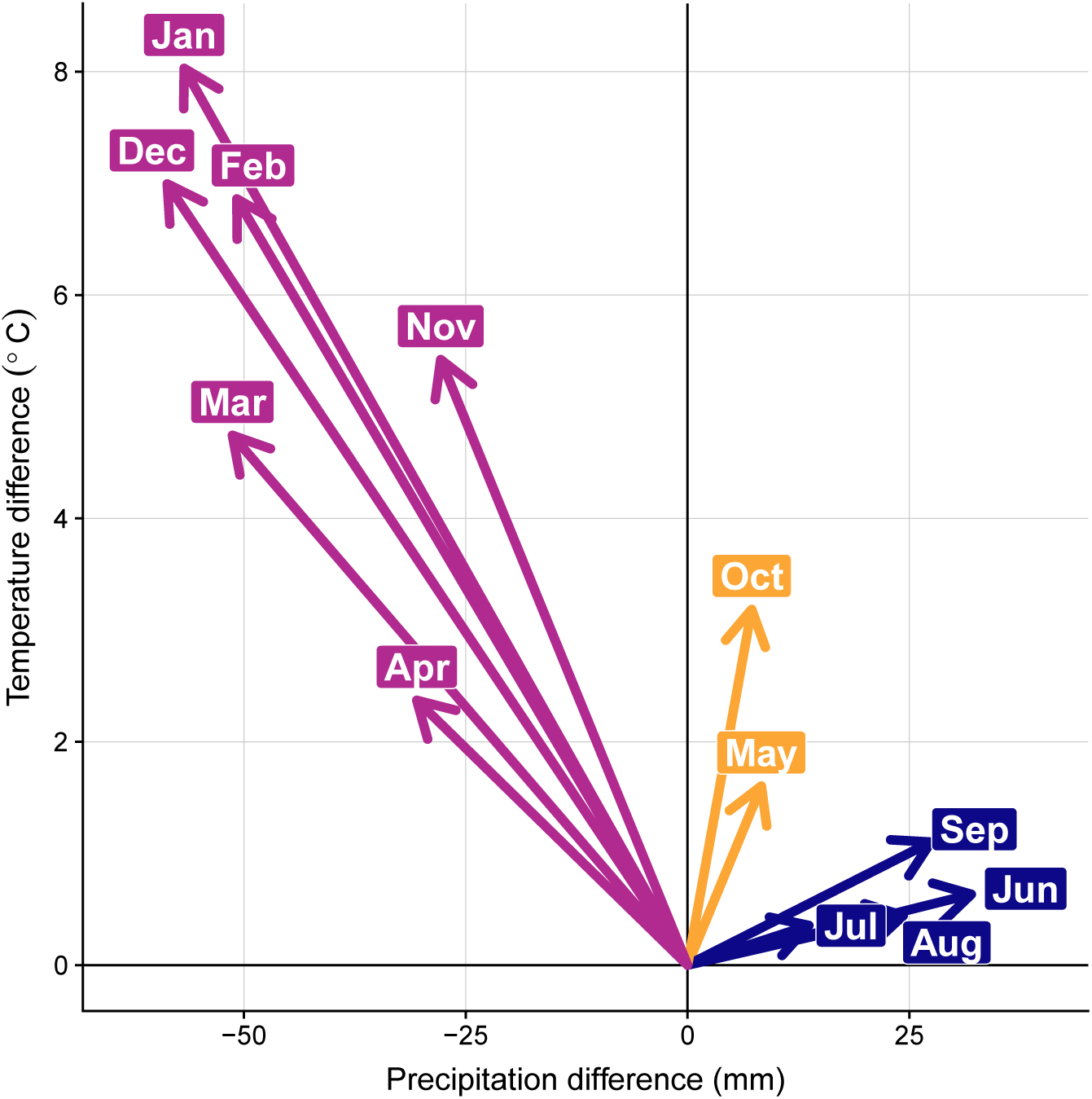
Seasonal variation in marginality, as measured by the difference between climatic conditions at wood stork GPS locations vs. the entire study area for each month. Each arrow represents selection in a given month for both precipitation (X-axis) and temperature (Y-axis): Wood storks select drier and warmer areas during the winter (November–April in fuchsia), wetter areas during the summer (June–September in dark blue), and intermediate conditions in transitional seasons (May and October in orange).

## Discussion

In this study, we investigated seasonal similarity in both geographic space (range) and ecological space (climatic niche) of the partially migratory wood storks in the southeastern U.S. Using home range metrics of overlap, we highlighted seasonal consistency in both spaces: a summer season (encompassing at least June–September, i.e., the wet season) and a winter season (encompassing at least November–February, i.e., the dry season) have been identified in both geographic and climatic spaces, punctuated by intermediate seasons (April–May and October). This resulted in wood storks using drier areas than generally available in the study area during the winter, but wetter areas during the summer, while using warmer areas year-round. However, higher overall similarity in the geographic than in the climatic space indicated the primacy of the former to lead to seasonal space use, with climate alone unable to fully explain temporal patterns.

The lowest value of geographic overlap occurred between the months of January and July/August, and is still greater than most values of climatic overlap. This can be in part explained by their migratory strategy: wood storks are partially migratory birds (Coulter et al. 1999, Picardi et al. 2020), which results in some individual migrating while others stay resident year-round. As a consequence, the overall (monthly) distributions at the population level are not as contrasted as would be for a fully migratory species, especially between migration phases. This result is highly similar to Golden Eagle (*Aquila chrysaetos*), which exhibited high overlap (BA = 87 ± 9) between breeding and non-breeding home ranges in western U.S. (Watson et al. 2014), but can also be compared with very low values (< 10) of overlap at the individual level for large migratory vertebrates, with minimum overlap as low as 3 (reindeer *Rangifer tarandus*) or 8 (red deer *Cervus elaphus*) documented in Norway (Cagnacci et al. 2016). Despite the very high geographic overlap in our study system, BA (Fieberg and Kochanny 2005) showed enough fluctuation to define clear seasons in the geographic space as well.

On the other hand, similarity in the climatic space was surprisingly low, and defined narrower seasons. While BA delineated clear seasons in the climatic space, their values were consistently lower than the corresponding ones in the geographic space, indicating higher fidelity in space than for climate. In other words, seasonal ranges (breeding and non-breeding) are sought after for different features than their climatic characteristics, suggesting that wood storks could be consider “niche switchers” (Nakazawa et al. 2004), reflecting seasonality in their needs during the breeding and non-breeding phases. Over the long term, birds have been shown to somewhat track climate change, however at a slower pace than the actual climatic shift (Devictor et al. 2008). Altogether this seems to indicate that birds’ fidelity to specific areas could counteract their need to track climatic conditions at both fine (within year, this study) and large (across years, Devictor et al. 2008) temporal scales. However, and contrary to our expectations, climatic similarity was greater during the wet season than during the dry season, as shown by larger BA values in between June–September than during the rest of the year. This is potentially a consequence of the low climatic variability during summer throughout the southeastern U.S.: it is wet and hot everywhere in the range, and despite wood storks occupying a larger area, available conditions are narrower than during the winter.

Selection of climatic conditions—in terms of temperature and precipitation—was remarkably stable within each season (especially during the summer), even though average climatic conditions still vary greatly from month to month in the study area. This is another indication of spatial constraint: instead of tracking climate conditions directly, wood storks settle in specific geographic areas, which directly defines and constrains the conditions available there in a given season. These areas are in turn associated with climatic conditions different than the rest of the range; these conditions however still vary between each season, in agreement with niche switching, whereby distinct ecological regimes are sought after in different seasons. Wood storks are heavily tied to prey availability during the breeding season, which is directly regulated by local hydrology of the wetland (Kahl 1964, Gawlik 2002). As a result, wood storks are potentially more sensitive to local variations in weather (especially rain) than average monthly climatic conditions (Borkhataria et al. 2012). As wood storks are constrained to their nests during the breeding season (generally February–May, Coulter et al. 1999), their winter range corresponds to the driest areas of the year-round distribution, which are associated with the best foraging opportunities (Coulter et al. 1987, Herring and Gawlik 2011).

Historically, South Florida was host to large breeding populations of wading birds, including wood storks, although their number have sharply declined with the drainage of South Florida’s wetlands (Frederick and Ogden 2003). The date of nest initiation has been delayed by several months (February–March vs. historical November–December initiations) in response to deteriorating habitat conditions (Frederick et al. 2009). With sufficient sample size (i.e., enough individuals monitored simultaneously and over several years to bring a good generalization power), our framework could be used to highlight fine-scale dynamics within single years, and investigate temporal trends (e.g., directly related to the timing of migration) over time. Wading birds (white ibises *Eudocimus albus* and wood storks), and specifically their timing of nesting, are considered an indicator of Everglades restoration success (Frederick et al. 2009), so understanding spatiotemporal dynamics for their migration is important to be able to detect changes from restoration vs. climatic conditions.

We used a simplified definition of the niche in two dimensions only—arguably, temperature and precipitation are amongst the most critical climatic variables that define niche dynamics (Luoto et al. 2007, Bucklin et al. 2015). However, the ecological space is highly dimensional, and by extension, so are ecological niches, even restricted to their climatic form. Our approach measuring overlap is by no means restricted to a plane. While measures of multi-dimensional overlap exist (see for instance an alternative approach of a multivariate index of niche overlap based on Tukey depth in Cerdeira et al. 2018), it is unclear how the resulting indices can be compared to a 2-dimensional index of geographic overlap, as overlap mathematically decreases with higher dimensionality. A first step is thus to reduce the ecological space to two dimensions, for instance focusing on the first two axes of a multivariate analysis, which typically account for most of the information (see e.g., Dallas et al. 2017), at the expense of a more difficult interpretation. In this particular case of temporally-varying niches, the K-select (Calenge et al. 2005) provides a natural way to find commonalities between monthly climatic niches, which are essentially similar to individual habitat selection with varying availability. Such an approach, given relevant data over the entire range, would allow to include other temporally-varying factors potentially critical to the dynamics of wood storks, like water levels, which are largely managed in South Florida (Gawlik 2002).

Finally, it can be noted that not all environments that are varying temporally are also seasonal—both short- and long-term changes are present in nature. Many arid regions are subject to unpredictable rainfall with no temporal correlation within the year, leading to nomadism (Jonzén et al. 2011, Teitelbaum and Mueller 2019). On the other extreme, global changes such as climate change (Harvey 2016) or land-use change (Song et al. 2018) potentially affect the entire planet, with direct impact on species ranges (Di Marco and Santini 2015). While we focused on seasonality in this manuscript, our framework is perfectly suited to handle less predictable movements, such as nomadism or range shifts. Furthermore, our approach does not rest on assumptions regarding movement strategies, nor is it constrained to considering yearly cycles as is the case for seasonal patterns. Given suitable data at the right temporal scale, investigating any temporal pattern will allow to highlight how the niche evolves in both geographic and ecological spaces, and identify either short- or long-term shifts.

While seasonality can be addressed in both spaces separately (home range dynamics in the geographic space, e.g., Lesage et al. 2000, dynamic habitat use in the ecological space, e.g., Basille et al. 2013), little attention has been devoted to a joint approach. At the individual level, Peters et al. (2017) present an elaborate approach to 1) identify migratory tactics in individual roe deer (*Capreolus capreolus*) in partially migrating populations, and 2) describe environmental factors which are different in both groups—and would explain the different tactics. We aimed to fill the gap at the population or species level by providing a simple, yet very general approach to explore seasonal variability in geographic and ecological spaces, which should prove very useful in dynamic ecosystem (La Sorte et al. 2018). For instance, the correspondence between the geographic and the ecological space is at the core of species distribution modeling, which infers potential range maps by considering all suitable areas, and predicts future potential ranges under changing conditions, such as climate change (Elith and Leathwick 2009). While in a stable, saturated system, ecological niches provide a direct equivalence to the geographic range (Soberón and Peterson 2005), it becomes increasingly critical to account for variation (seasonality, long-term trends) in natural ecosystems (Pearman et al. 2008).

## Supporting information

Media Supporting Information

Metadata S1: GIF version

Metadata S1: MP4 version

Metadata S1: MOV version

## Acknowledgments

We thank David Bucklin for extracting temperature and precipitation data from the Climate Research Unit dataset. Clément Calenge provided useful feedback on a previous version of this manuscript. Financial support for this work was provided by the U.S. Geological Survey’s Southeast Region, the U.S. Fish and Wildlife Service, the Army Corps of Engineers, the National Park Service, the Environmental Protection Agency (STAR Fellowship to R.B.), and the U.S.D.A. National Institute of Food and Agriculture (Hatch project 1009101). Handling of Wood Storks was conducted under U.S. Fish and Wildlife Service Threatened and Endangered Species Permits TE051552, TE801914, and TE37498B-0; Florida Fish and Wildlife Conservation Commission Special Purpose Permit WX07646; Georgia Department of Natural Resources Scientific Collecting Permit 29-WJH-05-79; U.S. Geological Survey Bird Banding Permit 22598-E; and University of Florida IACUC Protocol #E013 and ARC Protocol 002-11BGL. Any use of trade, firm, or product names is for descriptive purposes only and does not imply endorsement by the U.S. Government.

## Notes

#### Summary of Updates

Second (minor) revision of the paper.

https://basille.github.io/wost-seasonality/

