## Supplementary material for "Joint seasonality in geographic and ecological spaces, illustrated with a partially migratory bird": Media Supporting Information

MATHIEU BASILLE

*Department of Wildlife Ecology and Conservation, Fort Lauderdale Research and Education Center, University of Florida, Davie, FL 33314, USA.*

JAMES WATLING

*Department of Biology, John Carroll University, University Heights, OH 44118, USA.*

STEPHANIE ROMAÑACH

*Wetland and Aquatic Research Center, U.S. Geological Survey, Fort Lauderdale, FL 33314, USA.*

RENA BORKHATARIA

*Department of Wildlife Ecology and Conservation, Everglades Research and Education Center, University of Florida, Belle Glade, FL 33430, USA.*

Submitted to *Ecosphere* on: November 1, 2019

### Metadata S1: Wood Stork monthly ranges and climatic niches

In this animation (available as `Video_S1.gif`, `Video_S1.mp4` and `Video_S1.mov`), each panel presents wood stork monthly ranges (left and center) and climatic niche (right) during the year. Range maps show current GPS locations in a given month in black with kernel contour lines, and all locations from the entire dataset shown as a reference in gray, on a background of temperature (left, from blue for colder temperatures to red for warmer temperatures) or precipitations (center, from yellow for dry conditions to blue for wet conditions). Climatic niches show current conditions used in a given month (precipitation on the X-axis, and temperature on the Y-axis) in black, in comparison to conditions available in the study area in a given month in gray. The marginality arrow defines the difference between average available conditions and average used conditions.
