## Supplementary figures and images for "Joint seasonality in geographic and ecological spaces, illustrated with a partially migratory bird"

### Metadata S1: GIF version

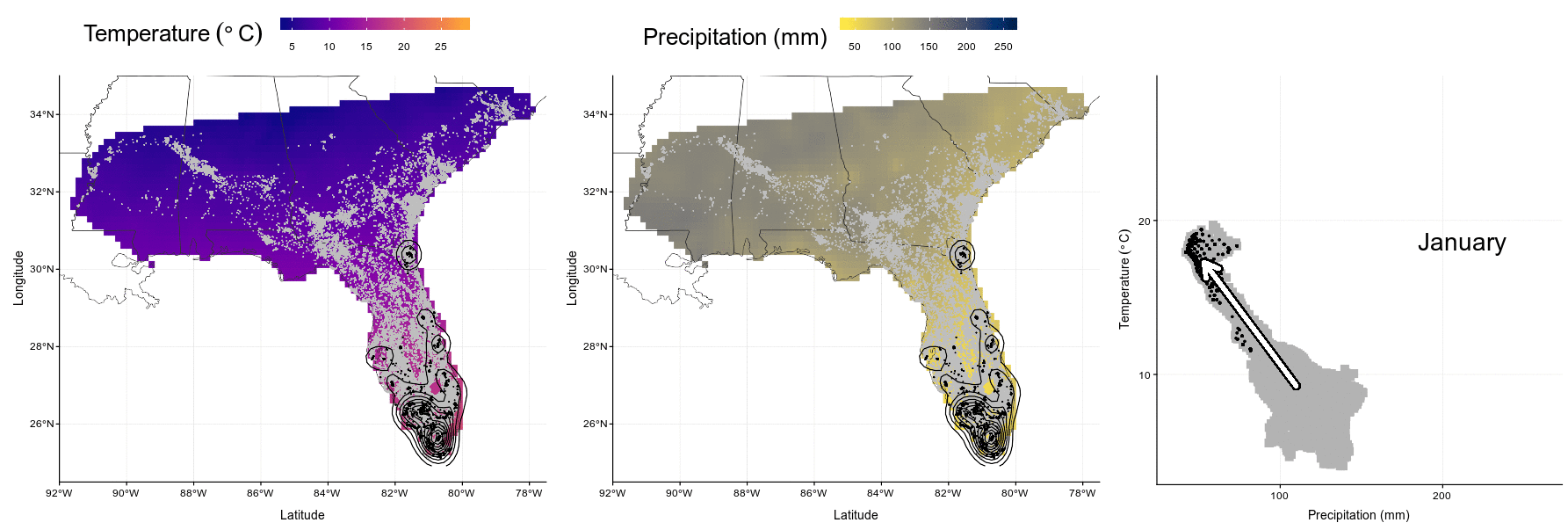
